# Quantitative Analysis of Interactive Behavior of Mitochondria and Lysosomes Using Structured Illumination Microscopy

**DOI:** 10.1101/445841

**Authors:** Qixin Chen, Xintian Shao, Mingang Hao, Zhiqi Tian, Chenran Wang, Fei Liu, Kai Zang, Fengshan Wang, Peixue Ling, Jun-Lin Guan, Jiajie Diao

**Affiliations:** School of Pharmaceutical Sciences, Shandong University, Jinan 250101, China; Department of Cancer Biology, University of Cincinnati College of Medicine, Cincinnati, OH 45267, USA; Shandong Academy of Pharmaceutical Science, Key Laboratory of Biopharmaceuticals, Engineering Laboratory of Polysaccharide Drugs, National–Local Joint Engineering Laboratory of Polysaccharide Drugs, Jinan 250101, China; Department of Biochemistry, University of Illinois at Urbana-Champaign, Urbana, IL 61801, USA

**Keywords:** mitochondria, lysosome, mitophagy, membrane fusion, super-resolution imaging

## Abstract

Super-resolution optical microscopy has extended the spatial resolution of cell biology from the cellular level to the nanoscale, enabling the observation of the interactive behavior of single mitochondria and lysosomes. Quantitative parametrization of interaction between mitochondria and lysosomes under super-resolution optical microscopy, however, is currently unavailable, which has severely limited our understanding of the molecular machinery underlying mitochondrial functionality. Here, we introduce an *M*-value to quantitatively investigate mitochondria and lysosome contact (MLC) and mitophagy under structured illumination microscopy. We found that the *M*-value for an MLC is typically less than 0.4, whereas in mitophagy it ranges from 0.5 to 1.0. This system permits further investigation of the detailed molecular mechanism governing the interactive behavior of mitochondria and lysosomes.

## Introduction

The crosstalk between mitochondria and lysosomes is involved in many cellular processes. For instance, mitophagy, a process that selectively removes redundant or damaged mitochondria, plays an important role in regulating the number of intracellular mitochondria and maintaining mitochondrial functions.^1^ Dysregulated 2 4 mitophagy is implicated in many diseases, such as neurodegenerative diseases and cancer.^2-4^ To date, mitophagy has been often studied at the cell level through methods such as flow cytometry,^5^ enzyme-linked immunosorbent assay,^6^ western-blot,^7^ and confocal fluorescence microscopy.^8-10^ These methods only report the cumulative level of mitophagy and ignore individual mitophagy events from the fusion between single mitochondria and lysosome pair. Although confocal fluorescence microscopy can detect mitophagy using mitophagy-specific dye, it is difficult to distinguish subcellular structures at a resolution beyond 200 nm.^11,12^ Moreover, confocal fluorescence microscopy does not provide details for the interactive behavior of individual mitochondria and lysosome pairs. Thus, a novel strategy is needed to capture detailed information on the crosstalk between individual mitochondria and lysosome pairs.

Recently developed super-resolution fluorescence microscopy techniques, such as stimulated emission deletion (STED)^,11,13,14^ structured illumination microscopy (SIM),^12,15,16^ and stochastic optical reconstruction microscopy (STORM),^17-19^ as well as other single-molecule super-resolution imaging techniques, ^20,21^ have provided new tools for investigating interactions between organelles at the subcellular level. And the interaction between mitochondria and lysosomes have been brought into focus using these techniques. For example, SIM super-resolution imaging has revealed a new type of interaction, mitochondria and lysosome contact (MLC). ^12,22^ Unfortunately, a parametrization system for the assessment of the interactive behaviors such as MLC and mitophagy under a super-resolution microscope is currently unavailable.

To fill this gap, in this toolbox paper, we propose an analysis system, *M*-value, to quantitatively analyze the crosstalk between mitochondria and lysosomes at the subcellular level. The *M*-value in MLC is less than 0.4, whereas the *M*-value in mitophagy ranges from 0.5 to 1.0. Thus, this *M*-value system provides a robust platform to quantitatively analyze the interactive behavior of subcellular organelles under super-resolution microscopy.

## Results

### MLC in living HeLa cells under SIM

To investigate MLC events in living HeLa cells using super-resolution imaging, we first incubated cells with a commercially available mitochondrial probe (Mito-Tracker Green, MTG, 100 nM) and a lysosomal probe (Lyso-Tracker Red, LTR, 200 nM) for 30 min, and then observed mitochondria (green) and lysosomes (red) under a SIM (Fig. 1A). The intracellular mitochondria showed spherical, rod-shaped or filamentous arrangement in an irregular manner at a resolution of approximately 150 ± 24.67 nm (*n*=10) (Fig. S1). Mitochondria are approximately 0.3 μm in width and 0.6-10.0 μ,m in length (Fig. S2), which is consistent with our previous results.^12^ In particular, there were more filamentous mitochondria than spherical and rod-shaped ones, a benchmark of healthy cells. In contrast, the diameter of spherical lysosomes (red color) was 0.6 ±0.16 μm (*n*=200) (Fig. S3). Both MTG and LTR allow for high-resolution staining under SIM, which enabled us to capture and understand the dynamic process of MLC (Fig. 1A, B). MLC events occur frequently in living cells (Fig. 1B) with a dynamic process from contact to separation (Fig. 1B, white arrows). In addition, we observed that mitochondria were surrounded by multiple lysosomes (Fig. 1C). Various types of MLC events, ^12,22^ such as point contact (Fig. 1E-7), extended contact (Fig. 1E-8), and surrounding contact (Fig. 1E-9) were also observed.

**Figure 1.**
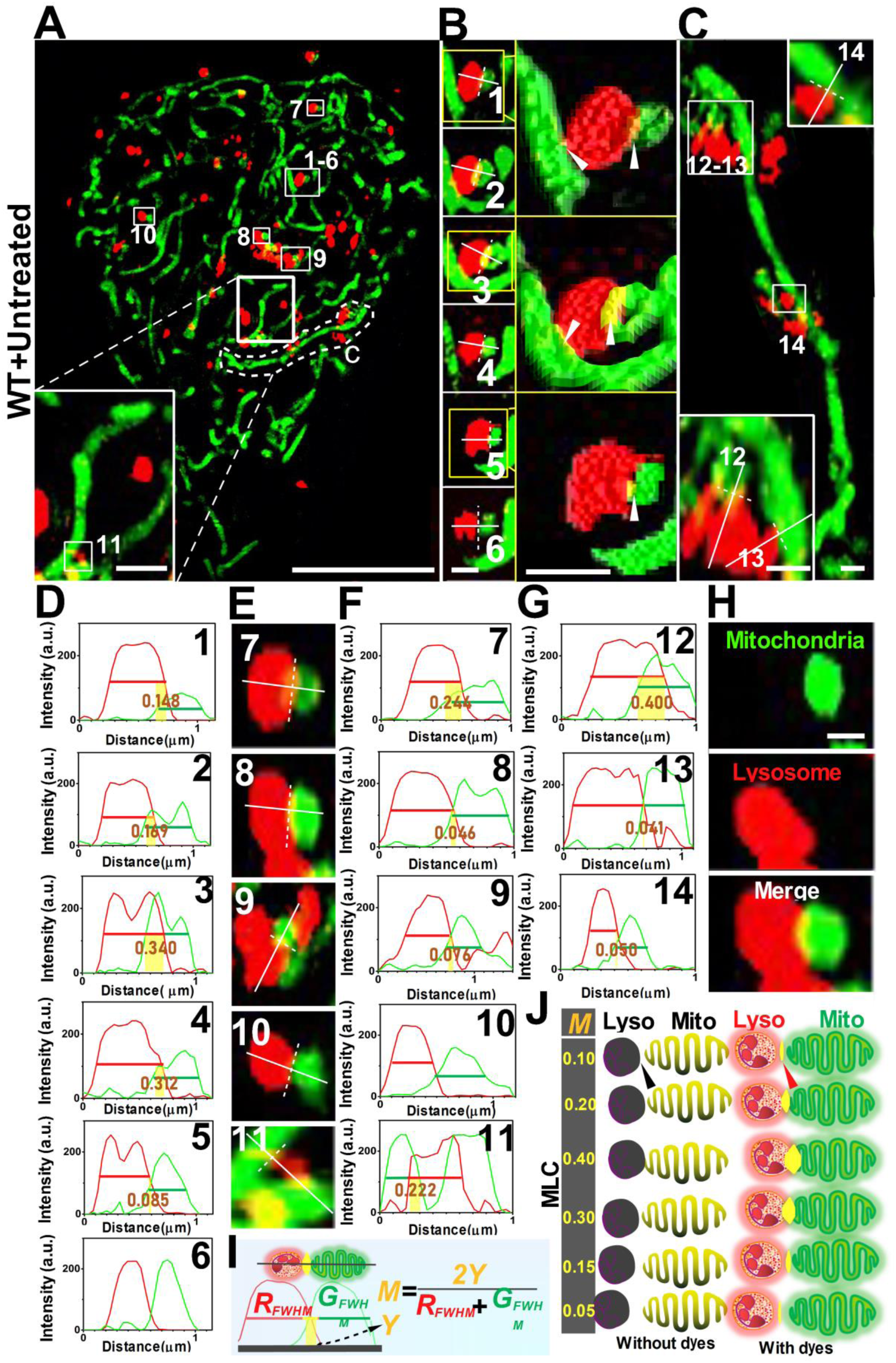
Whole cell quantitative analysis of mitochondrial-lysosome contact (MLC) in living cells. (**A**) SIM image of mitochondria (Green) and lysosomes (Red). 1-11 represent MLC events. (**B**) A representative dynamic process of MLC in a living HeLa cell. White solid lines indicate the fluorescence intensity shown in (**D**). (**C**) Partial enlargement of (**A**). White solid lines indicate the fluorescence intensity shown in (**G**). (**E**) and (**H**) Representative MLC events. White solid lines indicate the fluorescence intensity shown in (**F**). (**I**) Schematic diagram of calculation formula for the *M*-value of MLC. (**J**) *M*-value range of MLC dynamic events. Scale bars: **A** 5.0 μm, **B**, **C**, **E**, and **H** 0.5 μm, insets of **A** 0.5 μm.

### Quantitative analysis of whole-cell MLC

Recently, it has been reported that dysregulated contact sites formed by mitochondria and lysosomes are linked to Parkinson’s disease.^23^ We reason that proteins involved in MLC will provide a new perspective and drug targets for the treatment of this disease. A quantitative analysis platform to analyze MLC at the subcellular level would therefore allow us to understand the biological functions and evaluate the therapeutic effects of different drugs. Here, we introduce a quantitative analysis system, *M*-value that is derived from the full-width at half-maximum (FWHM) of organelle images, for understanding MLC events at the subcellular level. FWHM refers to the full width of the image at half-maximum value and can directly reflect the resolution of image. MTG-stained Mitochondria and LTR-stained lysosomes result in yellow spots when they merge (Fig.1H). Herein, we use the following calculation formula to define the *M*-value of the merge.

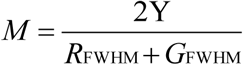

*R*_FWHM_ indicates the FWHM of red (lysosome); *G*_FWHM_ indicates the FWHM of green (mitochondria); Y indicates the merging distance between *R*_FWHM_ and *G*_FWHM_.

We will demonstrate that the *M*-value is a simple and well-defined number that can be used to quantitatively assess MLC events in live cells. We first observed that a typical MLC event (Fig. 1B, D) generates an ensemble of *M*-values below 0.4. We hypothesize that an *M*-value 0.4 can be the upper bound to characterize dynamic MLC events. To support this idea, we calculated the *M*-values of a large number of MLC events (*n*=100) (Fig.1C,E,F,G and Fig. S4-8) and found that the *M*-values of all MLC events were indeed below 0.4 (Fig. 1J).

### Quantitative analysis of whole-cell mitophagy

Mitophagy has been commonly studied by co-localizing fluorescently stained mitochondria and autophagosomes with confocal microscopy in cells.^24^ However, this strategy is not suitable for the quantitative analysis of mitochondria-lysosome interactions at the subcellular level. To address this issue, we set out to use the *M*-value to quantitatively analyze mitophagy.

Mitophagy was triggered by the incubation of HeLa cells with 10.0 μM carbonyl cyanide m-chloropheny hydrazone (CCCP), a common mitochondria damage inducer, for 12 h.^12,24,25^ Cells were then stained with MTG and LTR and observed under SIM (Fig. 2A). Treatment with CCCP broke mitochondria into spherical shapes with varying sizes compared to untreated cells, which primarily showed filamentous mitochondria (Fig. 1A). In addition, CCCP treatment induced more frequent mitophagy (Frame 1-3 in Fig. 2A), as evidenced by the appearance of more yellow spots resulting from fusion between mitochondria (green) and lysosomes (red) (Fig. 2A inset 1-3 and Fig. 2B). This result indicates that more mitophagy occurs under pathological conditions. Because dysfunctional mitophagy is closely associated with many diseases, ^2-4^ a quantitative analysis of mitophagy is of great significance for the drug development.

**Figure 2.**
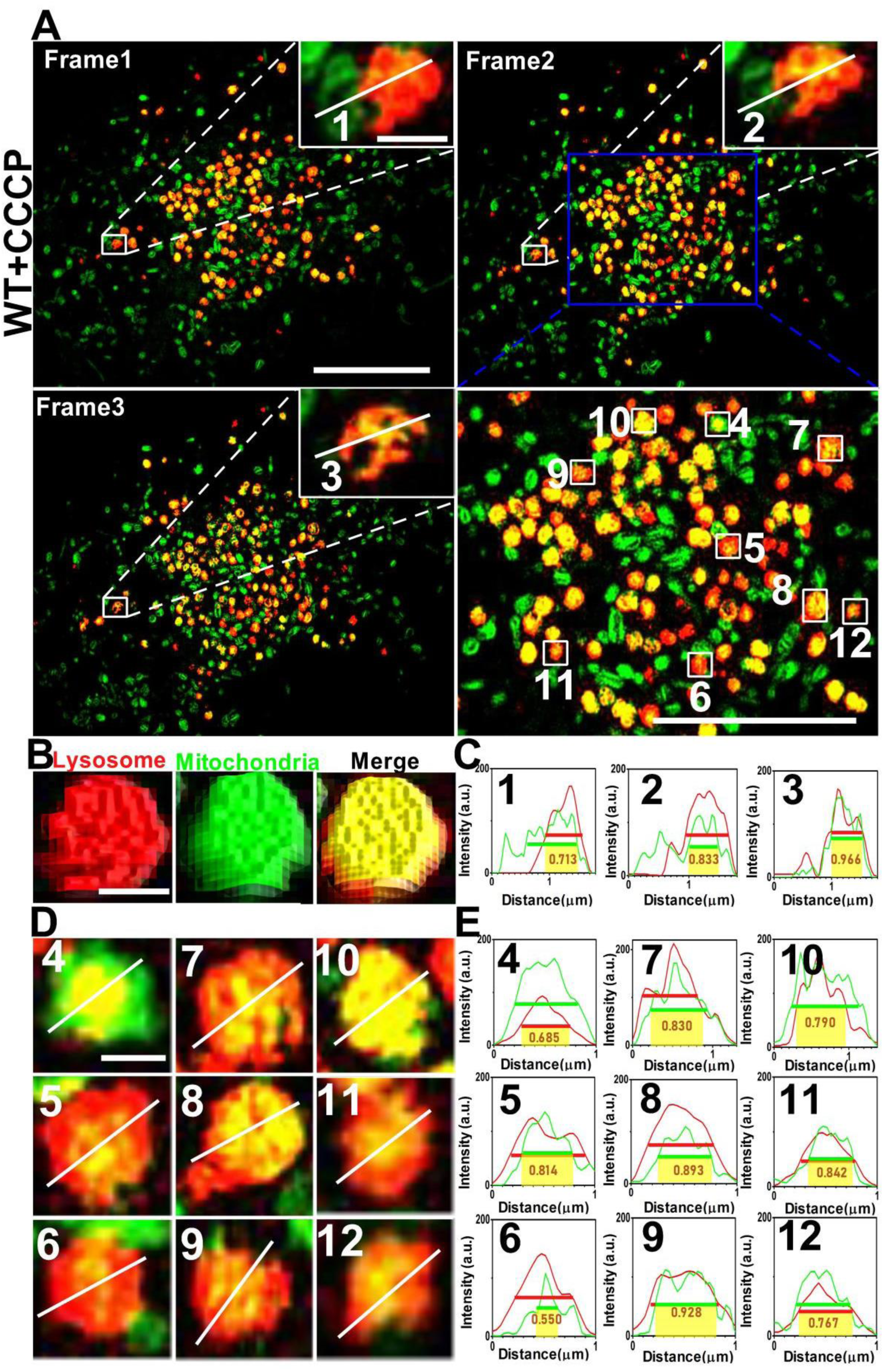
Quantitative analysis of fusion between mitochondria and lysosome in CCCP-treated cells. (**A**) Frame 1-3 of mitochondrial and lysosome fusion events. White solid lines indicate fluorescence intensity shown in (**C**). (**B**) 3D-SIM surface plot of representative fusion between mitochondria and lysosome. (**D**) Representative fusion events in (**A**). White solid lines indicate fluorescence intensity shown in (**E**). Scale bars: **A** 5.0 μm, **B** and **D** 0.5 μm, inset of **A** 0.5 μm.

With this goal in mind, we calculated the *M*-value of the mitophagy after the cell was exposed to CCCP for 12 h (Fig. 2A blue enlargement, and Fig. 2D). The *M*-value of the mitophagy event is greater than 0.5, and some even reaches close to 1.0 (Fig. 2C, E). To test whether the *M*-value can be applied to all mitophagy events, we performed a quantitative analysis to more mitophagy events (*n*=100) (Fig. S9-12). The *M*-values in all mitophagy events consistently range from 0.5 to 1.0 (Figure 3A). Thus, the *M*-value platform provides a robust system to differentiate MLC (0-0.4) and mitophagy (0.5-1.0) in physiological and pathological conditions.

**Figure 3.**
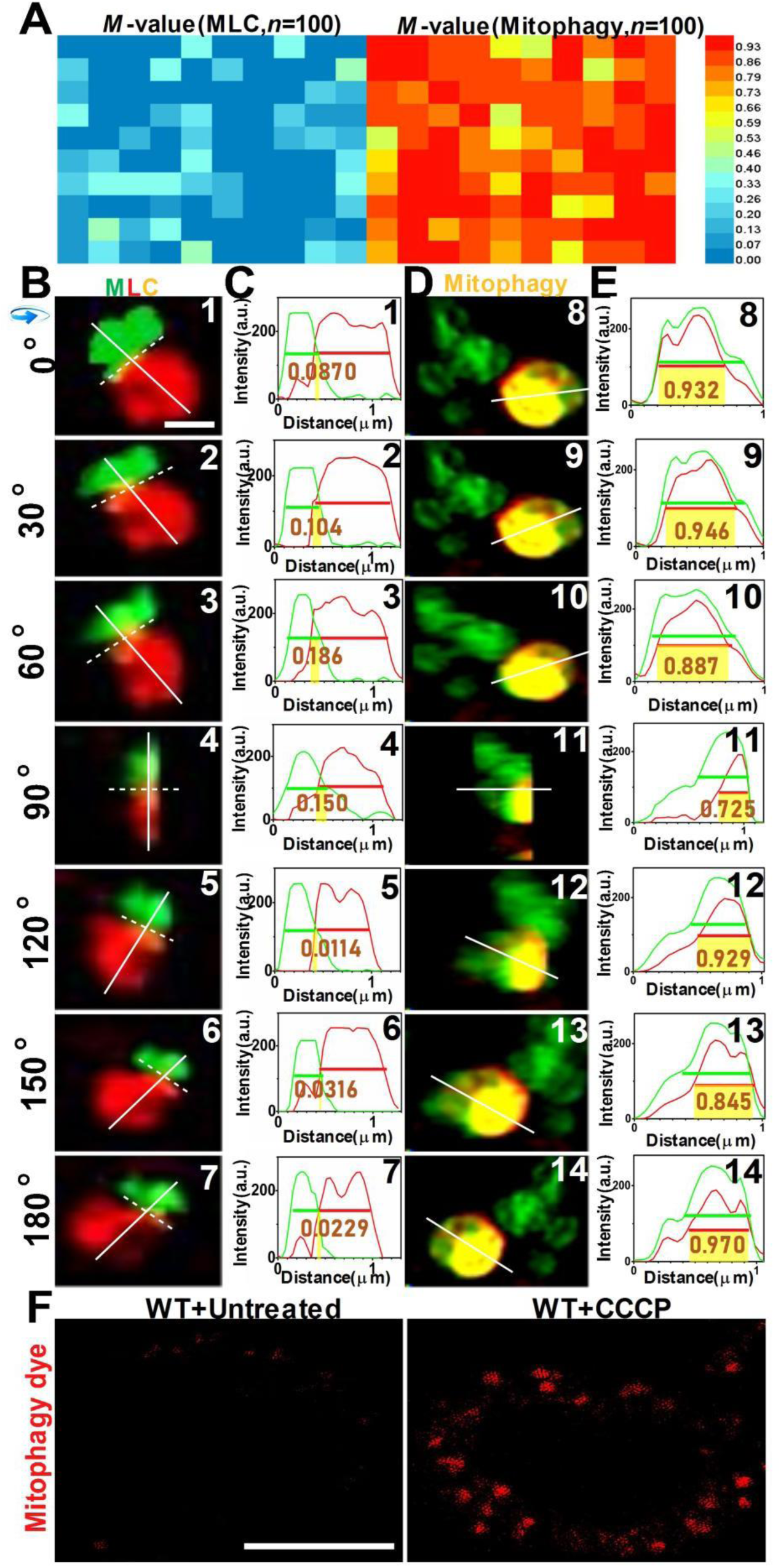
*M*-value of WT cells with and without CCCP treatment. (**A**) *M*-value distribution of 100 MLC events in normal cells and 100 mitophagy fusions in CCCP-treated cells. (**B**) A MLC event at different angles. The solid white line indicates the fluorescence intensity shown in (**C**). (**D**) A mitophagy at different angles. The white solid line indicates the fluorescence intensity shown in (**E**). (**F**) Cells incubated with mitophagy dye with or without CCCP treatment. Scale bars: **B** 0.5 μm, **F** 5.0 μm.

To verify the applicability of using the *M*-value for MLC and mitophagy, we next applied the *M*-value to quantitative analysis of an MLC (Fig. 3B, C) and a mitophagy event (Fig. 3D, E) at different rotation angles (0°-180°). We found that the *M*-value for MLC and mitophagy is consistent at different observation angles. Finally, CCCP-induced mitophagy was confirmed with a commercially available mitophagy detection dye, ^24^ which generates higher red fluorescence compared with untreated cells (Fig. 3F).

### Comparison of interactive behavior of mitochondrial and lysosome using epi-illumination fluorescence microscopy, confocal microscopy, and SIM

To demonstrate the advantages of SIM, we compared the performance of epi-illumination fluorescent microscopy, confocal microscopy, and SIM in resolving mitochondria-lysosome interaction using the same staining process with MTG and LTR (Fig. 4). Mitochondria and lysosomes appeared as green and red plaques in untreated cells under an epi-illumination fluorescence microscope, and a large area of yellow plaque was generated in the overlaid image (Fig. 4A). One cannot capture any information about MLC events at this spatial resolution, which may even cause a misinterpretation of mitophagy in untreated cells. Figure 4B showed the morphology of mitochondria and lysosomes under a confocal microscope. Although confocal microscopy successfully located mitochondria and lysosomes at the subcellular level, its spatial resolution (∼500 nm, Fig. S13) is not enough for the quantitative investigation of the detailed mitochondria-lysosome interaction below 100 nm, compared with SIM (Fig. 4C).

**Figure 4.**
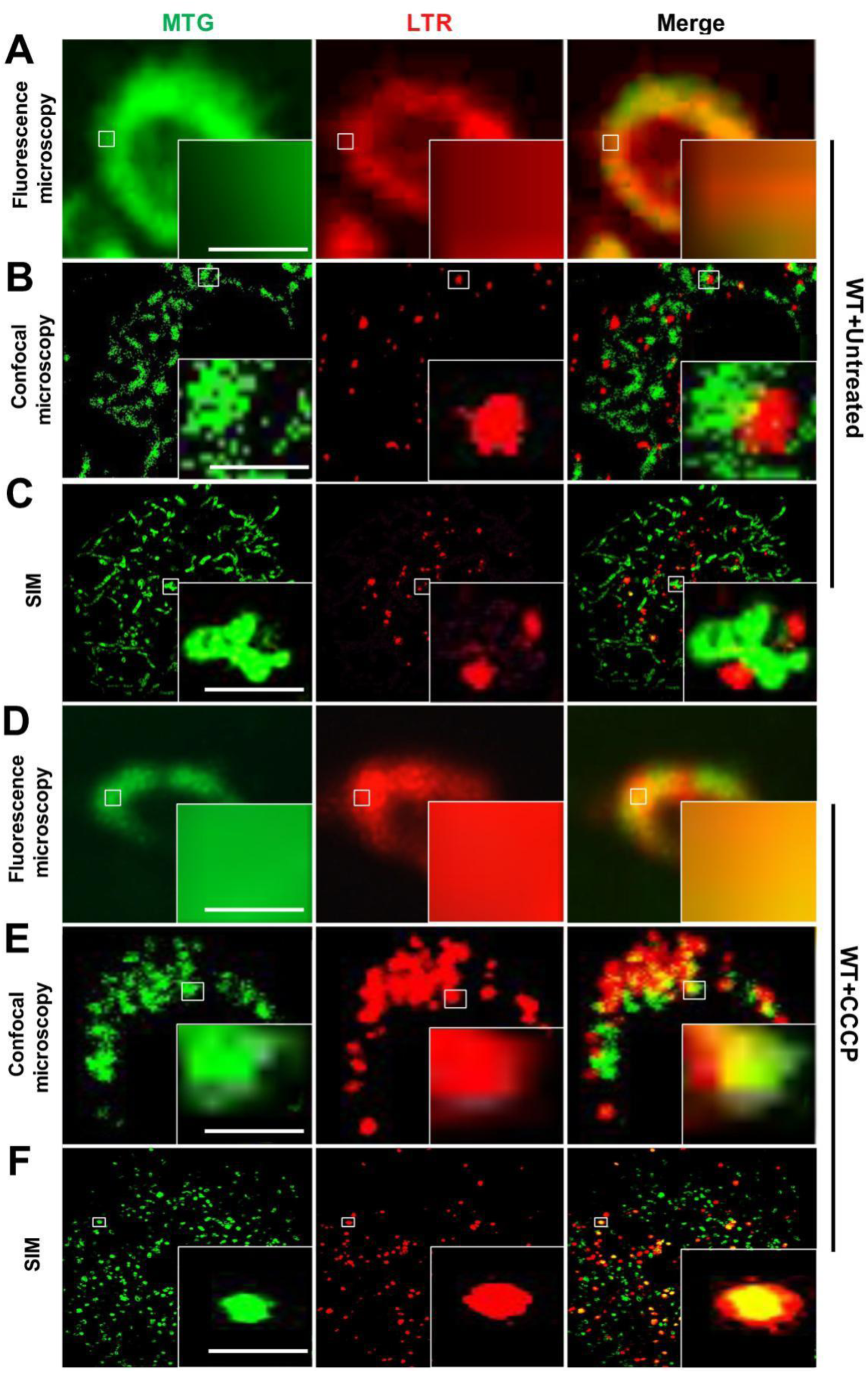
Comparison of interactive behavior of mitochondrial and lysosome using fluorescence microscopy, confocal microscopy and SIM. Mitochondrial and lysosome were stained with MTG and LTR. Images of untreated cells under fluorescence microscopy (**A**), confocal microscopy (**B**), and SIM (**C**). Images of CCCP-treated cells under fluorescence microscopy (**D**), confocal microscopy (**E**), and SIM (**F**). Scale bars: 1.0 μm.

In order to further investigate the mitochondria-lysosome interaction under pathological conditions using different optical microscopy techniques, we incubated 10.0 μM CCCP with HeLa cells for 12 h before staining with MTG and LTR. As expected, neither epi-illumination fluorescence microscopy (Fig. 4D) nor confocal microscopy (Fig. 4E) was able to clearly capture mitophagy at the subcellular level. In contrast, SIM provides excellent imaging quality that can resolve the mitochondria-lysosome interaction (Fig. 4F).

### Application of *M*-value to ATG13 and FIP200 knockout cell

Since the *M*-value system is feasible to quantitatively separate MLC (<0.4) and mitophagy (0.5-1.0), we then applied *M*-value to analyze the crosstalk of mitochondria and lysosomes in ATG13 knockout (ATG13 KO) and FIP200 knockout (FIP200 KO) HeLa cells. Both ATG13 and FIP200 play important roles in the process of autophagy and mitophagy (Fig 5A).^26,27,28,29^ The endogenous ATG13 and FIP200 in wild-type (WT) cells were knocked out with a CRISPR/Cas9 gene editing assay (Fig 5B). The ATG13 KO and FIP200 KO cells were then exposed to media with or without 10.0 μM CCCP for 12 h prior to staining with MTG and LTR. In untreated ATG13 KO cells (Fig 5C), mitochondria showed a continuous rod shape, and interacted with lysosomes as MLC (*M*-value, 0.245) (Fig 5E-1). After CCCP treatment, damaged mitochondria appeared as granules with various sizes (Fig 5C), which is similar to those in CCCP-treated WT cells (Fig 2A). MLC (*M*-value, 0.186) also occurred under CCCP treatment (Fig 5E-2). In contrast, compared to CCCP-treated WT cells (Fig 2A), mitophagy events in CCCP-treated ATG13 KO cells significantly decreased. In FIP200 KO cells, no mitophagy was observed before and after CCCP-treatment (Fig 5D, E-3, E-4), whereas MLC still occurred. These results demonstrated that ATG13 and FIP200 are required for mitophagy. The *M*-value system is applicable to distinguish the two events in autophagy-defective cells.

**Figure 5.**
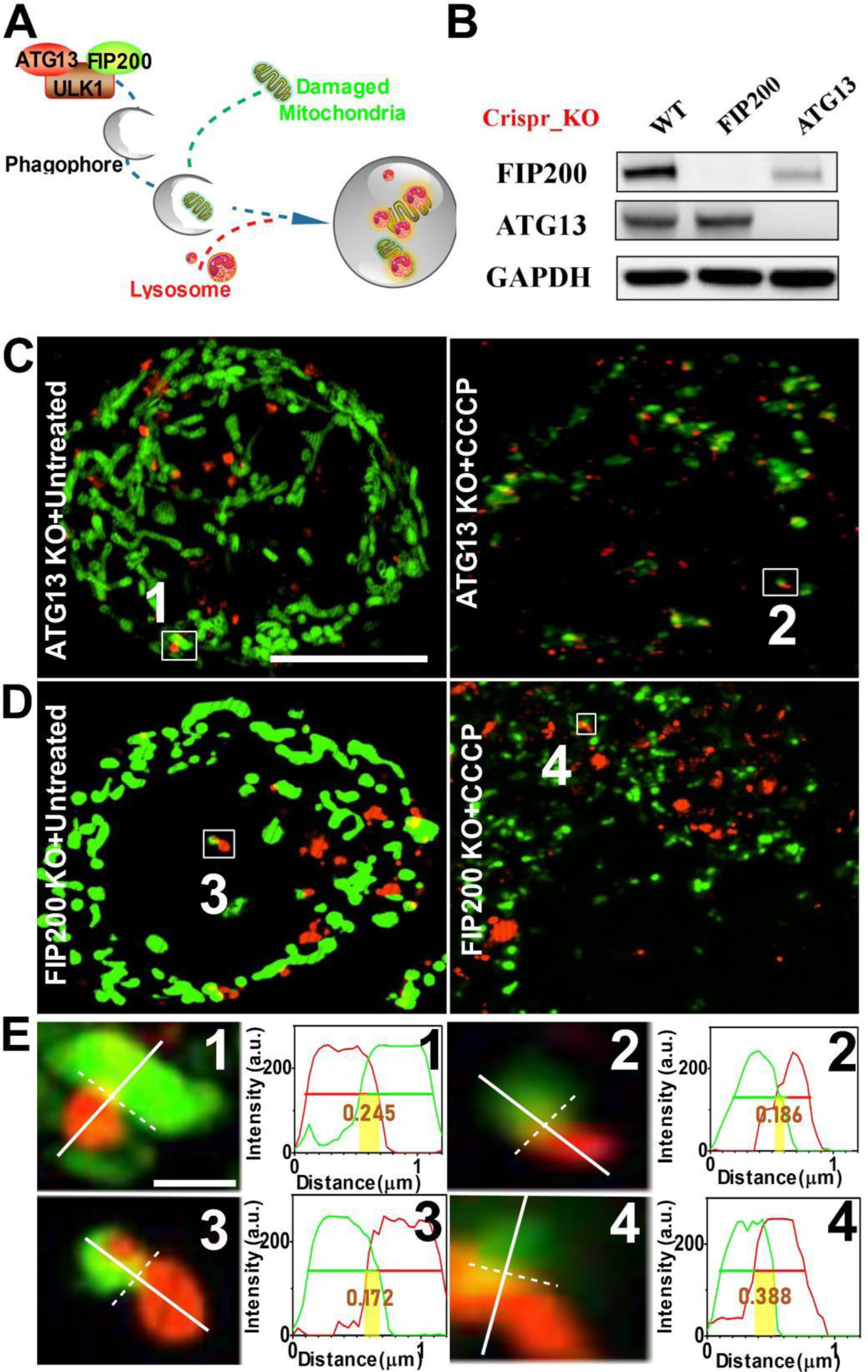
ATG13/FIP200 knockout cells under SIM. (**A**) Illustration of the role of ATG13 and FIP200 in mitophagy. (**B**) Western blot for detecting the protein expression of ATG13 and FIP200. Images of untreated and CCCP-treated ATG13 KO cells (**C**) and FIP200 KO cells (**D**). (**E**) Representative *M*-value calculations indicate MLC events in ATG13KO cells (**1,2**) and FIP200 KO cells (**3,4**) with or without CCCP treatment. Scale bars: **C** 5.0 μm, **E** 0.5

## Discussion

Mitochondria are important organelles for energy conversion in eukaryotic cells and are involved in a variety of biological processes such as intracellular homeostasis, proliferation, senescence, and death. ^30,31^ At the same time, functions of mitochondria are regulated by their interactions with lysosomes through processes such as MLC and mitophagy.^12,22,32,33^ MLC is widely found in cells, and it was recently discovered that RAB7 GTP hydrolysis plays a key role in the MLC process.^22^ Mitophagy is an autophagy process that selectively removes redundant or damaged mitochondria, which regulates the number of mitochondria in cells and maintains normal mitochondrial function.^2,4,34^ Herein, the interactive behaviors of individual mitochondria and lysosome pairs were observed under SIM. Compared to conventional tools such as epi-illumination and confocal fluorescence microscopy, SIM offers an improved spatial resolution to capture detailed information of the dynamic mitochondria-lysosome interactions in live cells.

Another bottleneck that limits our understanding of mitochondria-lysosome interaction is a lack of quantitative analysis platform for data generated from SIM. To address this challenge, we developed an *M*-value platform based on the FWHM of organelle images. This platform allows us to successfully differentiate MLC from mitophagy because their *M*-values are bounded within different range: an *M*-value below 0.4 implies MLC and an *M*-value ranging from 0.5 to 1.0 implies mitophagy. Thus, the *M*-value quantitative analysis system based on the super-resolution microscopy technique opens a new avenue to adopt automatic image analyzing software for high-throughput study of subcellular organelle interactions.

We envision that the *M*-value platform will specifically benefit high-throughput drug screening as an evaluation index of pharmacodynamics. Currently, high-throughput drug screening mainly relies on reporter gene systems, fluorescent labeling detection, micro-chemical technology, and fluorescence imaging technology. ^35^ Among them, the combination of fluorescence imaging and automatic analysis is an excellent screening strategy for micro-, sub-cellular image analysis in living cells. ^36^ However, the confocal microscopy systems commonly used in these high-throughput screening systems cannot provide sufficient spatial resolution to capture organelle interactions in live cells. ^37,38^ Combining artificial intelligence and SIM imaging with *M*-value system will significantly retrench the process of early drug discovery and shorten the drug screening cycle to promote the success rate of novel drug discovery.

## Materials and methods

### Materials

Mito-Tracker Green (#M7514, MTG) and Lyso-Tracker Red (#L12492, LTR) were obtained from Invitrogen (Eugene, Oregon, USA). Mitophagy dye was obtained from Dojindo Laboratories (#MD01-10, Kumamoto, Japan), Carbonyl cyanide 3-chlorophenylhydrazone (#045200, CCCP) was obtained from Thermo Fisherscientific (Grand Island, NY, USA). Penicillin-streptomycin (#15140163, 10,000 units/ml), Fetal bovine serum (#26140079, FBS), Dulbecco’s modified Eagle’s medium (#11965118, DMEM) and other cell culture reagents were obtained from Gibco BRL (Grand Island, NY, USA). Primary and secondary antibodies used in this study were GAPDH (#5174, Cell Signaling Technology, Beverly, MA, USA), FIP200 (#12436, Cell Signaling Technology, Beverly, MA, USA) and ATG13 (#13273, Cell Signaling Technology, Beverly, MA, USA) and HRP-linked anti-rabbit IgG (#7074, Cell Signaling Technology, Beverly, MA, USA). HeLa cells were a generous gift from Dr. Carolyn M. Price lab (University of Cincinnati).

### Cell culture

Cells were cultured in DMEM supplemented with 10% FBS, penicillin-streptomycin (100 units/ml) in a 5% CO_2_ humidified incubator at 37 °C.

### Live cell labeling

Cells were incubated with 100 nM MTG for 30 min and further co-incubated with 200 nM LTR at 37 °C for another 30 min in free DMEM, and washed with free DMEM 3 times and observed using a fluorescence microscopy, confocal laser scanning microscopy or OMX 3D-SIM super-resolution microscope.

### Confocal laser scanning microscopy

The images were obtained using a LSM-710 confocal laser scanning microscope (Carl Zeiss, Inc.) equipped with a 63×/1.49 numerical aperture oil-immersion objective. I1. and were analyzed with ZEN 2012 (Carl Zeiss, Inc.) and ImageJ software (National Institutes of Health). All fluorescence images were analyzed with ImageJ software (https://imagej.nih.gov/ij/).

### OMX 3D-SIM super-resolution microscope imaging and analysis

A total of 2×10^5^ cells were seeded on a glass bottom microwell dish and incubated with 2 ml of DMEM medium supplemented with 10% FBS for 24 h, followed by CCCP or vehicle treatment for 12h. After treatment, the cells were washed 3 times with pre-warmed free DMEM medium, stained with 100 nM MTG for 30 min, co-incubated with 200 nM LTR at 37°C for another 30 min, and washed with free DMEM for 3 times. Finally, cells were cultured in phenol-free medium (#1894117, Gibco, Grand Island, NY, USA) and observed under an OMX 3D-SIM super-resolution microscope (Bioptechs, Inc) that is equipped with an Olympus 100×/1.49 numerical aperture oil-immersion objective lens and solid-state lasers.

Firstly, the lasers (488 nm and 561 nm) were turned on and waited for 20 min to make the instrument thermally stable. After cleaning the lens, an immersion oil droplet (refractive index 1.516) was placed on the lens. Finally, the sample was placed on a tray for imaging. In order to reduce photobleaching, a 10 ms exposure time and 1% transmission excitation power was used to focus the sample and find the cells. Images were captured with an electron-multiplying charge coupled device (EMCCD) camera (Photometrics Cascade II) with a gain value of 3000 at 10 MHz. The exposure time was set to 50 ms per frame. Whole-frame images (512×512 pixels) were acquired using Z-stacks with a step size of 0.2 μm.

SIM images were analyzed with ImageJ software equipped a fluorescence intensity measurement tool. The intensity profile along the perpendicular (white solid lines, Fig. S14) of mitochondria-lysosome contact (white dash lines, Fig. S14) was used for the *M*-value calculation.

### CRISPR/Cas9-mediated knockout of FIP200 and ATG13 in Hela cells

The pX458 plasmid (pSpCas9(BB)-2A-GFP; Addgene) was used as the cloning backbone for expressing sgFIP200 and sgATG13. Two complementary oligos for each sgRNA were denatured, annealed and ligated into linearized pX458 vector digested by BbsI (New England Biolabs). Empty constructs and pooled pX458-sgRNA were transiently transfected into Hela cells respectively using Lipofectamine 3000 (Invitrogen). After 48 hr, the transfected cells were sorted based on the fluorescence of GFP (reporter) using a FACSAria cytometer (BD Biosciences). Sorted individual cells were cultured in a 96-well plate and were subjected to Western blot analyses. At least three different clones were pooled for functional experiments. The sgRNA sequences targeting FIP200 and ATG13 were based on the published literature using CRISPR/Cas9 library.^39,40^ sgRNA sequences of FIP200 and ATG13 are listed below:
sgFIP200: CAGGTGCTGGTGGTCAATGG
sgATG13-1: TCACCCTAGTTATAGCAAGA
sgATG13-2: CAGTCTGTTGTACACCGTGT
sgATG13-3: GACTGTCCAAGTGATTGTCC

### Western-blot assay

Cells were cultured in 3.5 cm diameter plates (80-90% confluence), washed by PBS buffer, and lysed for 15 min on ice using RIPA buffer (#C2978, Sigma, St. Louis, MO, USA) containing an anti-protease mix (#PI78415, Thermo Scientific, Waltham, MA, USA). Protein concentration was measured by BCA assay (#23225, Thermo Scientific, Waltham, MA, USA). Equal amounts of proteins were subjected to SDS-PAGE and immunoblotting as described previously.^41^

## Acknowledgements

This research was supported by 973 Program (No. 2015CB856304), National Institutes of Health (NIH, R35GM128837 to J.D.; R01NS094144 and R01CA211066 to J.G.; R01NS103981 to C.W.), Natural Science Foundation of Shandong Province (ZR2017PH072, ZR2017BH051, ZR2015QL007), and Key Research and Development Plan of Shandong Province (2018GSF121033). The Light Microscopy Imaging Center (LMIC) is supported in part with funds from Indiana University Office of the Vice Provost for Research. The OMX 3D-SIM microscpope was provided by NIH grant S10 RR028697.

## Author contributions

Q.C., X.S. and Z.T. collected all OMX 3D-SIM super-resolution microscope data. Q.C. and X.S. analyzed and processed the OMX 3D-SIM data. Q.C. cultured cell. M.H performed confocal laser scanning microscopy. K.Z., F.W., P.L., J.G., and J.D. conceived the project, designed the experiments, and wrote the manuscript with the help of all authors.

## Abbreviations

CCCP: carbonyl cyanide m-chlorophenyl hydrazone
FWHM: full-width at half-maximum
LTR: Lyso-Tracker Red
MLC: mitochondria and lysosome contact
MTG: Mito-Tracker Green
SIM: structured illumination microscopy

